# Long read subcellular fractionation and sequencing reveals the translational fate of full length mRNA isoforms during neuronal differentiation

**DOI:** 10.1101/2024.02.20.581280

**Authors:** Alexander J Ritter, Jolene M Draper, Chris Vollmers, Jeremy R Sanford

## Abstract

Alternative splicing (AS) alters the cis-regulatory landscape of mRNA isoforms leading to transcripts with distinct localization, stability and translational efficiency. To rigorously investigate mRNA isoform-specific ribosome association, we generated subcellular fractionation and sequencing (Frac-seq) libraries using both conventional short reads and long reads from human embryonic stem cells (ESC) and neural progenitor cells (NPC) derived from the same ESC. We performed *de novo* transcriptome assembly from high-confidence long reads from cytosolic, monosomal, light and heavy polyribosomal fractions and quantified their abundance using short reads from their respective subcellular fractions. Almost half of all transcripts exhibited association with particular subcellular fractions relative to the cytosol. Of the multi-isoform genes, 27% and 18% exhibited significant differential isoform sedimentation in ESC and NPC, respectively. Alternative promoter usage and internal exon skipping accounted for the majority of differences between isoforms from the same gene. Random forest classifiers implicated 3’ and 5’ untranslated region (UTR) GC-content and coding sequence (CDS) and UTR lengths as important determinants of isoform-specific sedimentation profiles. Taken together our data demonstrate that alternative mRNA processing within the CDS and UTRs impacts the translational control of mRNA isoforms during stem cell differentiation, and highlights the utility of using a novel long read sequencing-based method to study translational control.

## INTRODUCTION

Accurate eukaryotic gene expression requires messenger RNA (mRNA) assembly from precursor transcripts. Protein coding and regulatory sequences (exons) are distributed across expansive precursor messenger RNA transcripts. The spliceosome excises intervening non-coding sequences (introns) from pre-mRNA and ligates the exon sequences together generating a translation-competent mRNA (Y. Wang et al. 2015). Conserved sequences at exon-intron boundaries (splice sites) direct spliceosome assembly on each newly synthesized intron. Remarkably, the spliceosome can assemble different combinations of exon sequences to generate mRNA isoforms from a common pre-mRNA transcript (Konarska 1998; Wu et al. 1999). Alternative splicing (AS) not only generates isoforms with distinct protein coding potential, but also with different post-transcriptional regulatory capacity. For example, AS decisions that introduce premature termination codons induce nonsense mediated decay while other splicing events generate transcripts with distinct subcellular localization or translational control. In addition to generating alternative isoforms with distinct coding sequences (CDS), AS can produce isoforms that differ only in their untranslated regions (UTRs). Elements in the UTRs of mature mRNA play pivotal roles in post-transcriptional regulation. In the 5’ UTR, regulatory sequences like upstream open reading frames (uORFs) and internal ribosome entry sites (IRES) influence translation initiation efficiency (Hellen and Sarnow 2001; Morris and Geballe 2000; Weber et al. 2023). The 3’ UTR contains various elements such as microRNA binding sites and RNA-binding protein (RBP) recognition sites that modulate mRNA stability, localization, and translation (Ciolli Mattioli et al. 2019; Mayya and Duchaine 2019). Regulatory elements in the CDS can also influence the fate of mRNAs. For example the RBP, HuR, stabilizes target mRNAs by binding to AU-rich elements (AREs) within the CDS, preventing their degradation. Conversely, RBPs like TTP can promote mRNA degradation by binding to AREs in coding regions, leading to mRNA decay (Otsuka et al. 2019). Proteins like IGF2BP1 can bind to coding region instability determinants in the CDS of target mRNAs to enhance their stability (Weidensdorfer et al. 2009). By and large, AS confers complex and multidimensional consequences to the fate of mRNAs through shaping the *cis*-regulatory landscape of alternative isoforms (Castle et al. 2008; Pan et al. 2008; E. T. Wang et al. 2008).

Importantly, there is poor correlation between steady-state mRNA and protein levels in eukaryotic systems (Lian et al. 2001; Griffin et al. 2002; Cox, Kislinger, and Emili 2005; Schmidt et al. 2007; Fu et al. 2009). And while factors like mRNA stability and translation initiation efficiency play a role in this disparity, the substantial influence of AS on translational control is often overlooked. A number of methods exist to study translational control, which is the regulatory mechanism in eukaryotic cells that governs the efficiency and timing of protein synthesis from mRNA. A widely used method called Ribo-seq offers genome-wide insights into ribosome occupancy and translation dynamics by capturing single nucleotide-resolution ribosome footprints but can be vulnerable to artifacts and signal biases (Ingolia et al. 2009). RNC-seq captures ribosome nascent-chain complex-bound mRNAs to characterize the translatome, but it doesn’t provide ribosome footprints or ribosome density information (T. Wang et al. 2013). TRAP-seq utilizes epitope-tagged ribosomes to enable cell type-specific translation profiling, which generates data similar to RNC-seq which can be modified to produce ribosome footprints, but it relies on transgenic models and may not fully replicate endogenous ribosome behavior (Reynoso et al. 2015; Heiman et al. 2014). Frac-seq, which our proposed method builds on, isolates actively translating ribosomes and assesses translation efficiency by stratifying transcripts by the number of ribosomes they are associated with (Sterne-Weiler et al. 2013). However, it has the potential for selective bias toward highly abundant transcripts and it lacks single-nucleotide resolution of ribosome positions on mRNA. While each method has its strengths and weaknesses, one shared disadvantage is that they all involve the sequencing of short mRNA fragments from ribosome-protected or ribosome-associated mRNAs.

Short read RNA-sequencing methods struggle to accurately capture the complete structures of complex RNA isoforms (Steijger et al. 2013). In contrast, long read RNA-sequencing provides full-length reads that span entire transcripts, enabling precise characterization of intricate isoforms and annotation-agnostic detection of novel structures. The primary shortcoming of long read sequencing is its relatively lower throughput compared to short read sequencing platforms, limiting the depth of coverage for a given budget. To address this limitation and to maximize the benefits of both long read and short read methods, we employed a complementary approach. Here we introduce the development of long read Frac-seq to obtain full-length, ribosome-associated transcript isoforms with intact records of ribosome association, structural variation, and long-range interactions. We complement this data with short read Frac-seq to compensate for the loss of throughput and to provide a more complete and accurate representation of the translated transcriptome.

## RESULTS

### Characterization and measurement of a transcriptome generated solely from full-length transcripts captured in the model system

To investigate the relationship between alternative pre-mRNA splicing and isoform-specific mRNA translation we capitalized on the capability of long read sequencing to capture complete transcript structures of polyribosome-associated mRNA, without sacrificing throughput by generating both long read and short read Frac-seq libraries (Sterne-Weiler et al. 2013). We used human embryonic stem cells and neural progenitor cells (ESC and NPC, respectively) as a model system to characterize the translated transcriptome during early neuronal differentiation. The resulting samples were the cytosol, monosome, light polyribosome (2-4 ribosomes), and heavy polyribosome (≥5 ribosomes) fractions (Figure 1A). By utilizing the R2C2 method (Volden et al. 2018), our long read libraries, with mean read length 2 Kb and mean library size 500 K, were reinforced with improved base calling accuracy (93%) and with high-confidence transcript starts and ends. Fractionation of the long reads was employed to enhance the likelihood of detecting transcripts which may be preferentially associated with distinct polyribosome fractions. All long read libraries were pooled for *de novo* transcriptome assembly using Mandalorion (Volden et al. 2023), followed by rigorous quality control, filtering, and functional annotation using all three modules of the Functional IsoTranscriptomics analysis suite (de la Fuente et al. 2020).

**Figure 1.**
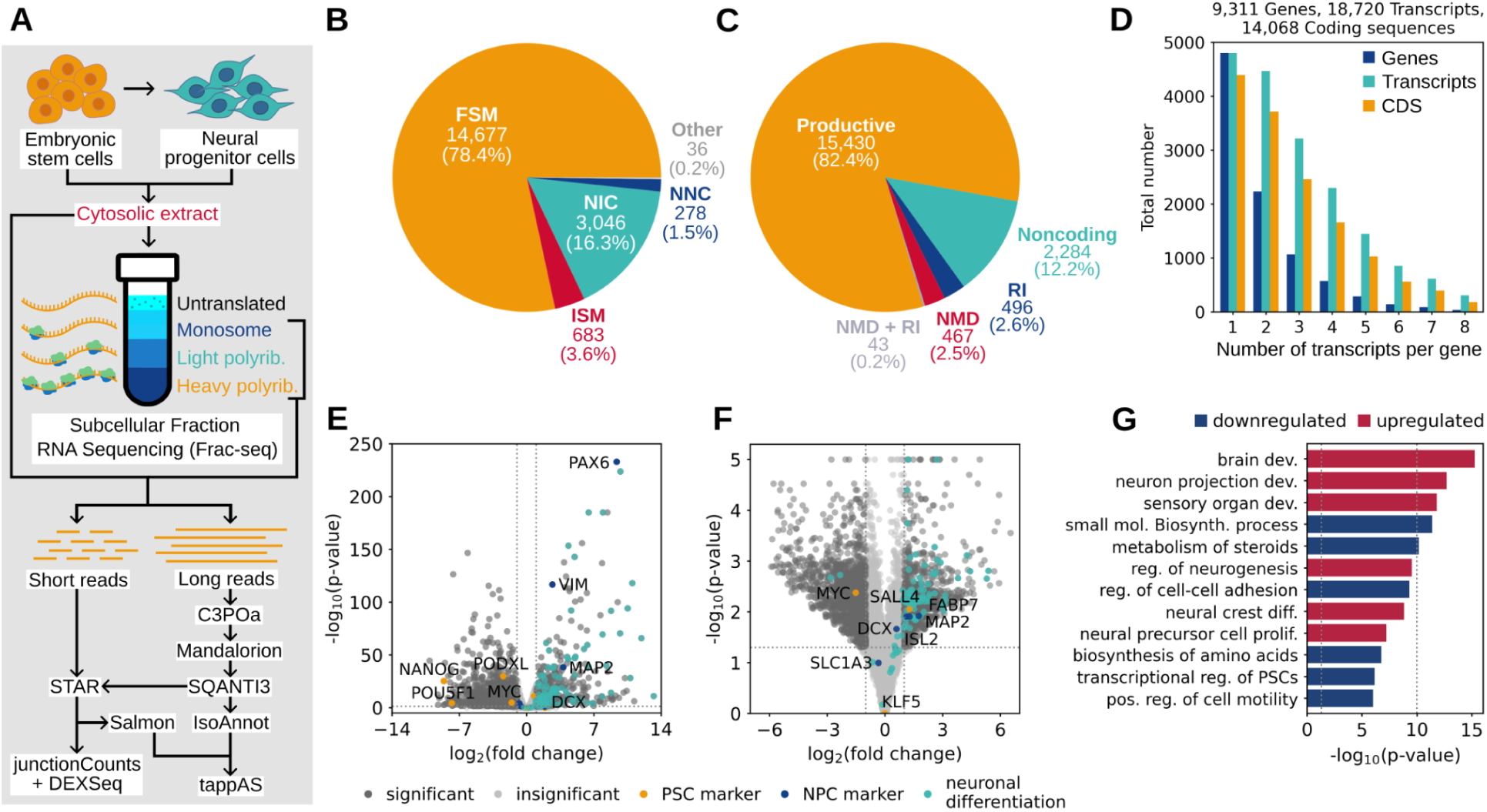
Experimental overview and characterization of the long read-derived transcriptome and the cytosol. (A) Schematic of the experiment and subsequent bioinformatic analysis workflow of the resulting cytosolic extract and fractionated, ribosome-associated short and long reads from ESC and NPC. (B) The transcriptome classified by SQANTI3-defined structural categories of spliced transcripts, including: Full Splice Match (FSM), Incomplete Splice Match (ISM), Novel in Catalog (NIC), Novel Not in Catalog (NNC) and intergenic or fusion transcripts (Other). FSM and ISM transcripts match annotated splice sites and junctions (in GRCh38.p13 Release 41), while NIC transcripts comprise novel combinations of annotated splice sites and junctions and NNC transcripts contain at least one unannotated splice site. (C) The transcriptome classified by productivity based on the detection of complete or incomplete open reading frames (productive or noncoding, respectively), premature stop codons (NMD) and retained introns (RI). (D) Stratification of the transcriptome by the number of isoforms and unique coding sequences per gene. (E-F) Gene-level (E) and transcript-level (F) differential expression between NPC and ESC cytosolic fractions. (G) Top 12 enriched Metascape pathways in differentially expressed genes between NPC and ESC cytosolic fractions.

The long read-derived transcriptome (LR transcriptome) consisted of 18,720 transcripts with 14,068 unique coding sequences (CDS), arising from 9,311 genes (Figure 1D). Of these, 1,274 and 784 transcripts, and 423 and 175 genes were unique to ESC or NPC, respectively. Transcripts were organized into SQANTI3-defined structural categories based on their fidelity to transcript structures in the GRCh38.p13 Release 41 primary assembly annotation (Harrow et al. 2012). 82% of transcripts matched the annotation, 17.8% were considered novel (containing either novel combinations of known splice sites and junctions or at least one novel splice site), and less than 0.2% were categorized as either genic or fusions (Figure 1B). Additionally, transcripts were categorized based on their productivity. We define productive transcripts as those encoding a full-length, canonical protein. Unproductive classes include: noncoding (lacking a complete open reading frame), nonsense-mediated decay (NMD) and retained intron (RI). 82.4% of transcripts were considered productive, while 12.2% were noncoding and the remaining 5.3% were NMD or RI (Figure 1C).

We used Salmon (Patro et al. 2017) to pseudoalign the fractionated short reads, with an average library size of 71.5 M reads, to the LR transcriptome; producing transcript-level quantification across the gradient. Using the cytosolic fraction, which represents the raw output of the nucleus, we next tested the baseline transcriptomic differences in NPC relative to ESC at the gene-level (Figure 1E) and at the transcript-level (Figure 1F) to reveal upregulation of NPC and neuronal differentiation markers and downregulation of pluripotency markers. Metascape (Zhou et al. 2019) pathways further encapsulated these observations (Figure 1G). Taken together, these results present the framework for an approach to integrate fractionated long and short reads to study translational control at isoform-level resolution.

### A large portion of transcripts exhibit distinct association with particular subcellular fractions

To discover if mRNA transcripts have distinct ribosome association profiles we clustered transcript-level expression trajectories across the gradient using tappAS (de la Fuente et al. 2020), revealing subpopulations of transcripts with clear enrichment in one subcellular fraction over the others (Figure 2A). Transcripts with significant enrichment (log_2_FC ≥ 1.0, p-value ≤ 0.05) in a particular subcellular fraction relative to the cytosol were designated, without mutual exclusivity, as associated with that fraction (Figure 2B). In fact, 46% and 39% of transcripts were significantly associated with a subcellular fraction in ESC and NPC, respectively. Subpopulations of transcripts enriched in particular subcellular fractions were extremely dissimilar (Jaccard similarity ≤ 0.08) across fractions within each cell type, with the largest Jaccard similarity of 0.18 and 0.19 between the heavy polyribosome fraction and the cytosol in ESC and NPC, respectively (Supplemental Figure 1). When stratified by productivity, monosome- and light polyribosome-associated transcript subpopulations exhibited pronounced incorporation of unproductive transcripts relative to the cytosol, while heavy polyribosome-associated transcript subpopulations displayed the opposite characteristic (Figure 2C). These findings support the hypothesis that levels of ribosome association may correlate with levels of translation. 5846 (64%) and 4972 (56%) genes in ESC and NPC, respectively, contained at least one subcellular fraction-associated – or differentially sedimenting (relative to the cytosol) – isoform. TMEM59 is an example of a gene with four differentially sedimenting isoforms in ESC (Figure 2D). Interestingly, endogenous post-transcriptional silencing of TMEM59 by miR-351 in neural stem cells has been implicated to promote neuronal differentiation (Li et al. 2012), although all four isoforms share the same 3’ UTR.

**Figure 2.**
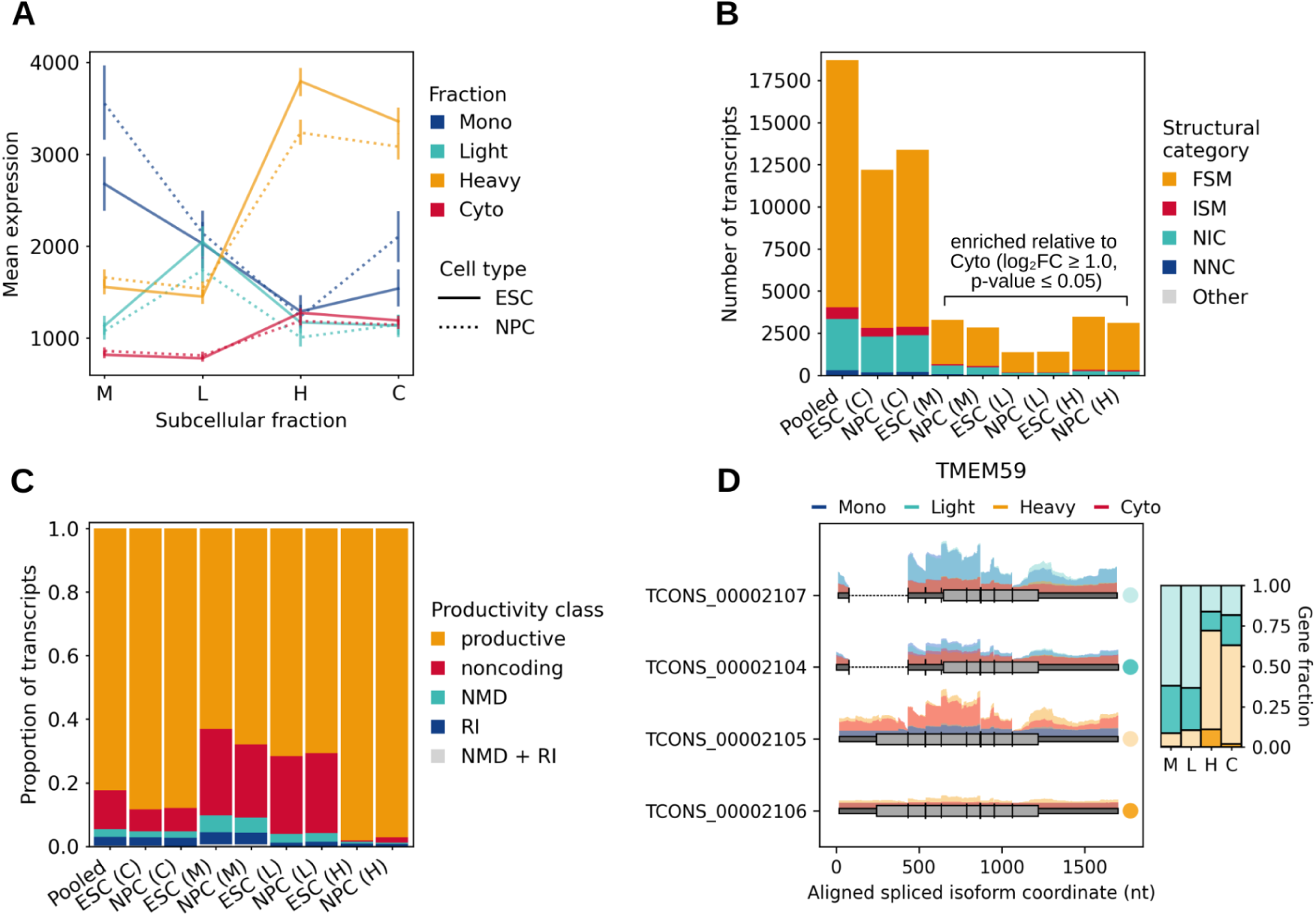
Establishing transcript ribosome association profiles. (**A**) Clustering of transcripts by their expression trajectories across the gradient; monosome (M), light polyribosome (L), heavy polyribosome (H) and the cytosol (C). (**B**) Extraction of fraction-associated transcripts based on significant enrichment (log_2_FC ≥ 1.0, p-value ≤ 0.05) in the monosome, light or heavy polyribosome fractions relative to their abundance in the cytosol. (**C**) Categorization of fraction-associated transcripts by productivity. (**D**) Differential sedimentation of 4 fraction-associated isoforms in TMEM59. Above spliced isoform models, histograms of short read support at exons are colored by their magnitude in each fraction. The stacked barplot summarizes the proportion of total gene expression each isoform contributes in each fraction.

### Alternative splicing confers functional consequences to the stability and translation of mRNAs

Because a substantial portion of transcripts were observed to have distinct ribosome association profiles, we postulated that alternative mRNA isoforms may likewise sediment discretely. To test this hypothesis, we calculated the expression of individual isoforms relative to all isoforms from the same gene within a given subcellular fraction. We found 1195 (27%) and 757 (18%) multi-isoform genes exhibiting differential isoform sedimentation in ESC and NPC, respectively. These transcripts reveal instances where alternative splicing could drive isoform-specific translational control. Changes in gene fraction across the gradient followed concordant patterns between ESC and NPC, suggesting that a given isoform in one cell type will likely associate with the same subcellular fraction in the other cell type (Figure 3A). In fact, only 102 transcripts in 75 genes exhibit divergent patterns of isoform sedimentation (Supplemental Figure 2). Among genes displaying differential isoform sedimentation, pathways in RNA processing, metabolism and splicing, along with DNA damage response were enriched in both ESC and NPC (Figure 3B). Pathways related to cell cycle checkpoints were unique to ESC, and pathways in noncoding RNA processing and metabolism were unique to NPC. Of the few genes demonstrating divergent isoform sedimentation across cell types, protein ubiquitination and endocytosis-related pathways were enriched.

**Figure 3.**
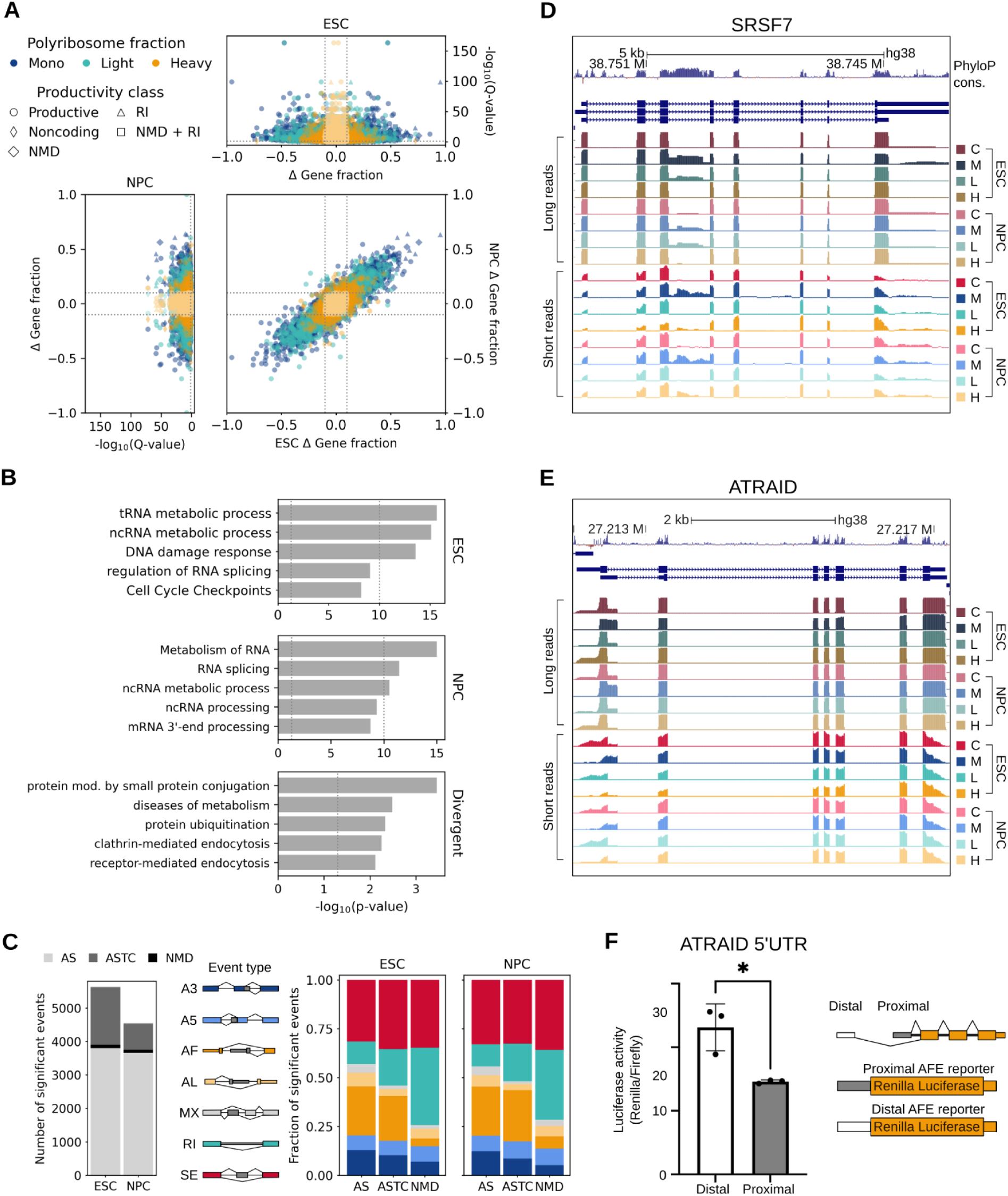
Differential isoform sedimentation across the gradient and functional outcomes of alternative splicing. (**A**) Volcano plots representing differential isoform sedimentation by changes in isoform gene fraction relative to the cytosol. 1209 and 757 genes, in ESC and NPC respectively, exhibit significant differential isoform sedimentation (|Δ Gene fraction| ≥ 0.1, Q-value ≤ 0.05). The central plot shows changes in isoform gene fraction, with Q-value ≤ 0.05, of isoforms present in both ESC and NPC. (**B**) The first two bar plots show a subset of enriched Metascape pathways in genes containing significant instances of differential isoform sedimentation in ESC and NPC. The third bar plot, labeled “Divergent”, depicts enriched Metascape pathways in genes displaying contrasting patterns of differential isoform sedimentation between ESC and NPC. (**C**) The first stacked bar plot categorizes significant alternative splicing events (|ΔΨ| ≥ 0.1, adjusted p-value or Q-value ≤ 0.05) as: alternative splicing (AS), alternative splicing coupled with translational control (ASTC), meaning splicing events that are differentially included across the gradient, and alternative splicing coupled with nonsense-mediated decay (NMD). The following two bar plots show the breakdown of event types comprising each category in ESC and NPC. (**D-E**) UCSC Genome Browser snapshot of long read and short read coverage at (**D**) SRSF7, exhibiting fraction-associated inclusion of a conserved retained intron, and at (**E**) ATRAID, exhibiting fraction-associated alternative first exon usage. (**F**) Luciferase assay measuring the translational impact of using either the distal or the proximal ATRAID 5’ UTR in HEK-293 cells.

To examine the types of alternative splicing (AS) that give rise to the diversity of the LR transcriptome, we categorized AS events as: AS (0.1 ≤ Ψ ≤ 0.9, adjusted p-value ≤ 0.05 within condition), ASTC (|ΔΨ| ≥ 0.1, Q-value ≤ 0.05 across subcellular fractions), or AS coupled with NMD (events that introduce a premature termination codon – PTC – and adhere to the mentioned cutoffs for significance) (Figure 3C). Notably, 36% and 19% of significant AS events were classified as ASTC in ESC and NPC, respectively. We found that alternative first exon, retained intron, and skipped exon events feature most prominently among ASTC events, while skipped exons and retained introns comprise the majority of NMD events. In our dataset, SRSF7 presents one complete and two partial retained intron events associated with NMD via induction of a PTC in the highly conserved SRSF7 intron 3 locus, which has been previously described to contain a conserved poison exon (Lareau et al. 2007; Königs et al. 2020) (Figure 3D). Preferential association of PTC-containing isoforms with the monosome, and modestly with the light polyribosome fraction, is consistent with our understanding of NMD’s effect on translation (Maquat et al. 1981; Nickless, Bailis, and You 2017; Celik et al. 2017). ATRAID (also known as APR3), a relatively poorly understood gene implicated to play roles in all-*trans* retinoic acid-induced apoptosis, osteoblast differentiation and some cancer types (Zhang et al. 2023; Ding et al. 2015), demonstrates marked patterns of alternative first exon usage across the gradient. The proximal first exon of ATRAID was preferentially spliced into the monosome-associated isoform, which may indicate its reduced translation. Indeed, a Renilla-firefly luciferase assay comparing Renilla incorporating either the proximal or the distal 5’ UTR of ATRAID in HEK293 cells exhibited significant differences in fluorescence (Figure 3F). Interestingly, the two isoforms of ATRAID may be functionally different, as the distal first exon contains an upstream open reading frame which may encode a signal peptide with potential importance to its localization with lysosomes (Ding et al. 2015). Collectively, these results demonstrate the widespread functional impacts of alternative splicing to the cytosolic fate of mRNAs.

### Intrinsic features and *cis*-elements correlate with transcript polyribosome profiles

Given that AS defines the *cis*-regulatory landscape of mature mRNAs, we hypothesized that intrinsic transcript features may encode the underlying regulatory grammar of ASTC. We define intrinsic transcript features as measurements and functional elements that are native to the sequence of a spliced transcript. To extract the relative weight of features on ASTC, we employed random forest classifiers (RFC) to perform feature selection. Across the LR transcriptome, we measured the length and GC-content of the transcript, CDS, and UTRs. Additionally, 5’ and 3’ UTR motifs, miR sites, uORFs, repeat sequences, AU-rich elements, coding capacity, the presence of a PTC and retained introns were included in the feature set. Given these features, RFCs were assigned binary classification tasks to predict the correct subcellular fraction for transcripts between every combination of subcellular fraction-associated transcript subpopulations at an 80:20 train:test split using 300 estimators. From the results, we extracted permutation feature importance, with 50 repeats, and found that the length of the CDS and UTRs, and the GC-content of the UTRs were important features for our models to correctly classify transcript polyribosome profiles.

Our findings that CDS and 3’ UTR length positively correlate with association with heavier polyribosome fractions is consistent with previous reports (Floor and Doudna 2016) (Figure 4A). This is not to be confused with ribosome density, which other groups have shown to be inversely correlated with CDS length (Zhao et al. 2017; Arava et al. 2003). Although a longer CDS can theoretically accommodate a greater number of ribosomes, the increased potential for incorporation of non-optimal or rare codons may trigger codon usage-dependent negative impacts to translation initiation and elongation (Lyu et al. 2021). Additionally, longer CDS and transcript lengths have been observed to be negatively correlated with translation initiation rates in the context of intrapolysomal ribosome reinitiation (Rogers et al. 2017). En masse, inference of ribosome association based on the CDS alone is likely too simplistic to make accurate predictions. To more clearly understand changes in feature length that may impact ribosome association, we also measured the change in CDS, 5’ UTR and 3’ UTR length relative to the dominant cytosolic isoform among isoforms belonging to genes with differentially sedimenting isoforms (termed gene-linked isoforms). Distinctly, monosome- and light polyribosome-associated isoforms displayed a clear signal of relatively shorter CDS and longer UTRs, while heavy polyribosome-associated isoforms remained largely similar or equivalent to the dominant cytosolic isoform (Figure 4B). Relatively longer 3’ UTRs in gene-linked isoforms are connected to strong effects on ribosome association, and isoforms with 5’ UTRs ≥ 1000 nt in length have been observed to be relatively poorly ribosome-associated relative to their shorter 5’ UTR-containing counterparts within the same gene (Floor and Doudna 2016). These phenomena could be due, in part, to potential increased inclusion of *cis*-regulatory elements in UTRs including miRNA target sites, uORFs and iron-responsive elements which can negatively impact mRNA stability and translation. Upon measuring LR transcriptome-wide GC-content, we found that GC-content of the 3’ UTR most clearly positively correlates with ribosome association (Figure 4C). This observation connects handily with findings that relate lower 3’ UTR GC-content with increased association with P-bodies and enhanced susceptibility to miRNA targeting (Courel et al. 2019). Among gene-linked isoforms, GC-content was decreased in the 5’ UTR and increased in the 3’ UTR in monosome and light polyribosome-associated isoforms (Figure 4D). The finding that lowly ribosome-associated isoforms have relatively higher 3’ UTR GC-content is concordant with reports that suggest an inverse relationship between 3’ UTR GC-content and mRNA stability (Litterman et al. 2019).

**Figure 4.**
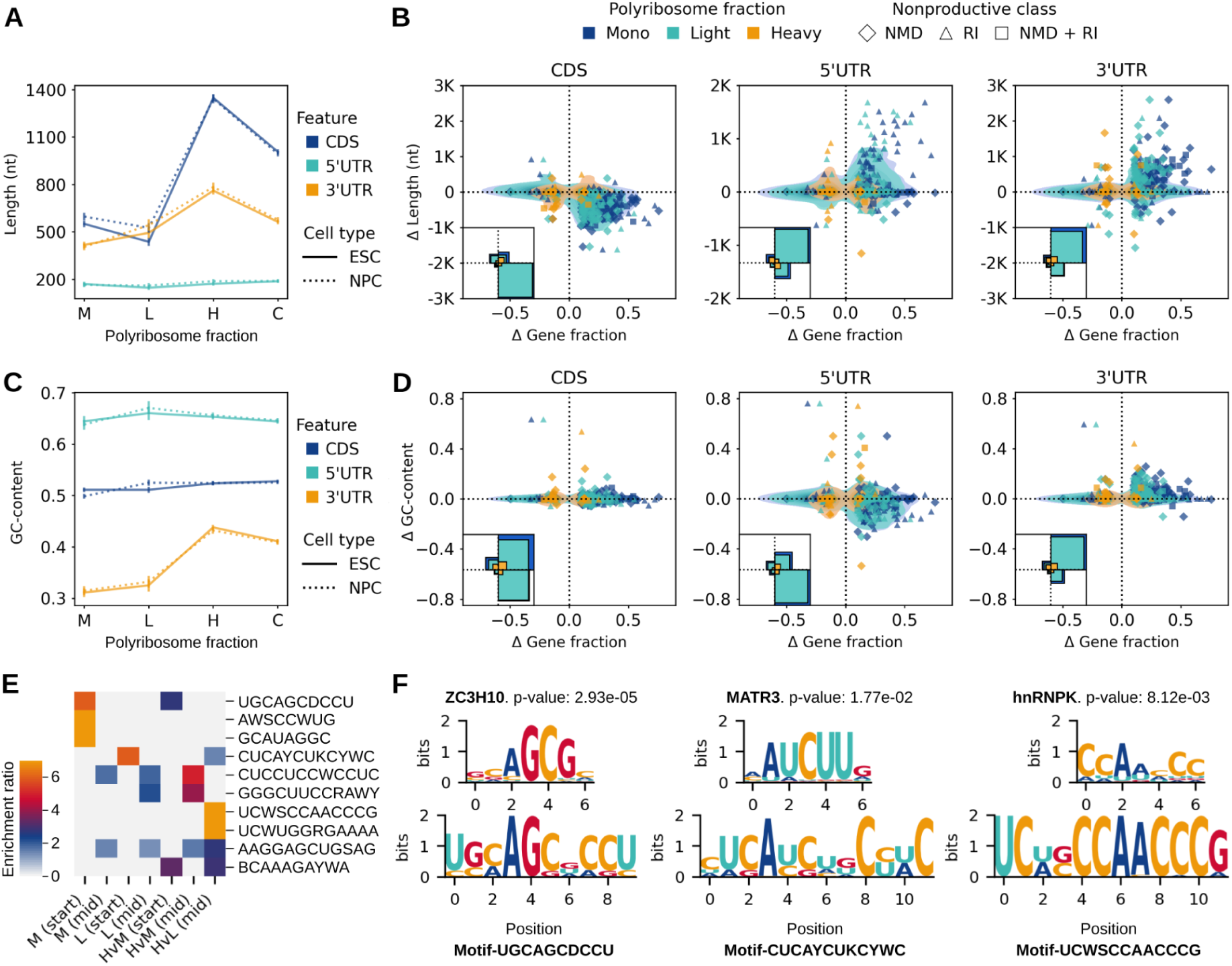
Analysis of features correlated with ribosome association profiles. (**A**) Mean length, with 90% confidence interval, of the CDS, 5’ UTR and 3’ UTR of fraction-associated coding transcripts. (**B**) Measurement of the change in isoform gene fraction relative to the cytosol and differences in CDS, 5’ UTR and 3’ UTR length of differentially sedimenting isoforms relative to the dominant isoform in the cytosol. Kernel densities for all coding isoforms are drawn with a 0.1 threshold, and NMD and RI isoforms are represented as scattered points. Subplots in the bottom left of each plot summarize the relative abundance of observations in each quadrant of their respective main plot, colored by fraction. (**C**) Mean GC-content, with 90% confidence interval, of the CDS, 5’ UTR and 3’ UTR of fraction-associated coding transcripts. (**D**) Measurement of the change in isoform gene fraction relative to the cytosol and differences in CDS, 5’ UTR and 3’ UTR GC-content of differentially sedimenting isoforms relative to the dominant isoform in the cytosol. Subplots, kernel densities, and scattered points were drawn with identical specifications to those in (**B**). (**E**) Enrichment of the top 10 most enriched HOMER-derived *de novo* RNA sequence motifs in skipped exons enriched in fractions. Fraction-enriched skipped exons were sliced and segregated into the first 30 nt (start), the last 30 nt (end) and the rest of the 30 nt windows comprising the exon body (mid). Enrichment ratio represents the relative enrichment of a given motif in the primary set of sequences versus the background (30 nt windows processed as mentioned, but from skipped exons not enriched in a given fraction). (**F**) Three examples of CISBP-RNA RNA binding protein sequence motifs with significant matches (p-value ≤ 0.05) to motifs identified in fraction-enriched skipped exons. ZC3H10 matches monosome-associated Motif-UGCAGCDCCU, MATR matches light polyribosome-associated Motif-CUCAYCUKCYWC and hnRNPK matches heavy polyribosome-associated Motif-UCWSCCAACCCG.

In addition to measuring length, GC-content and functional elements, we identified sequence motifs that are associated with particular subcellular fractions. To do this, we took fraction-associated skipped exons (FASE) that were determined to be significantly ASTC (ΔΨ ≥ 0.1, Q-value ≤ 0.05 across subcellular fractions) relative to the cytosol, sliced them into 30 nt windows, and separated them into three groups: start (first window), end (last window), and middle (all windows composing the exon body; excluding the first and last windows) to detect potentially exon start- or end-associated motifs. Each set of windows was complemented with a background set consisting of windows made from skipped exons that were not significantly ASTC in their given subcellular fraction. The heavy polyribosome fraction was tested for ASTC against the monosome and light polyribosome fraction to produce the heavy polyribosome set due to a dearth of FASEs relative to the cytosol. The resulting sets of FASEs included 600 exons on average for each subcellular fraction. Using HOMER (Heinz et al. 2010) on each set of windows, we found 133 motifs that were significantly enriched (p-value ≤ 0.05, FDR ≤ 0.2) in the target sequences over the background. Using SEA (Bailey and Grant 2021), we found that 63 of these motifs were enriched (p-value ≤ 0.05, enrichment ratio ≥ 1.1) in at least one set of FASE windows (Figure 4E). We used Tomtom (Gupta et al. 2007) to identify RBPs in the Ray 2013 Homo sapiens dataset (Ray et al. 2013) whose binding specificities match an enriched motif and selected the best-matched RBP (p-value ≤ 0.05). Three examples of RBPs with significant matches to enriched motifs are ZC3H10, MATR3, and hnRNPK which align to a monosome-, light polyribosome- and heavy polyribosome-associated motif, respectively (Figure 4F). We acknowledge, however, that ZC3H10 and MATR3 are largely uncharacterized and that RBP binding specificities are often multivalent and difficult to predict. Nonetheless, we report the presence of statistically significant sequence motifs enriched in FASEs. As a whole, these results suggest that ribosome association is impacted by the composition of intrinsic transcript features; likely with combinatorial effects.

## DISCUSSION

Here, we report the first integration of long read RNA sequencing with a translatomic method, which we call LR Frac-seq, and we describe an approach to integrate long read and short read Frac-seq to characterize the translated transcriptome in human ESC and NPC. We took a complementary approach to capitalize on the major strength of long read sequencing in capturing complete transcript structures, while leveraging short read sequencing’s significantly higher throughput for accurate quantification. Many examples of hybrid sequencing approaches have previously been applied to complex biological problems by other groups (Reese et al. 2023; Puglia et al. 2020). For the long reads, we employed the R2C2 method to generate high-confidence consensus sequences with high basecalling accuracy and well-defined transcript start and end sites. From these, we performed *de novo* transcriptome assembly to generate the set of full-length transcripts detected in the system, deemed the LR transcriptome. The much deeper fractionated short read libraries were utilized to quantify the LR transcriptome across the gradient, consisting of: the cytosol, monosome, light polyribosome (2-4 ribosomes), and heavy polyribosome (≥ 5 ribosomes) fractions. Indeed, the LR transcriptome does not comprehensively capture the entirety of the expressed transcriptome in ESC and NPC, as indicated by short read transcript-level mapping rates, but for this demonstration we chose to focus on the set of transcripts directly captured by LR Frac-seq, which were subsequently subjected to stringent quality control and filtration using SQANTI3 (de la Fuente et al. 2020). Despite the limited depth of the LR Frac-seq libraries, the resulting LR transcriptome still constituted a sufficient diversity of transcripts to encapsulate transcriptomic changes indicative of neural differentiation across cell types (Figure 1E and F) and isoform-level differences between subcellular fractions (Figure 3A). Highlighting one of the major benefits of long read sequencing, we found 3,324 transcripts with either novel combinations of known splice sites or ≥ 1 novel splice sites; accounting for 17.8% of the LR transcriptome.

We compared transcript abundances in subcellular fractions to their cognate cytosolic fractions to identify transcripts with enrichment in particular fractions relative to the cytosol, which represents the raw output of the nucleus. We found that nearly half of the LR transcriptome preferentially associates with subcellular fractions and that the proportion of productive transcripts associating with a given fraction directly correlates with ribosome association (Figure 5). Among multi-isoform genes expressed in both ESC and NPC, patterns of gene-linked differences in isoform sedimentation relative to the cytosol were largely congruent, suggesting that intrinsic transcript features may play a bigger role in ribosome association than cell type differences within our model. Using random forest classifiers to select features at the transcript level, we found that CDS, 5’ UTR and 3’ UTR length along with 5’ UTR and 3’ UTR GC-content were important features for the accurate prediction of transcript polyribosome profiles in our dataset. We then measured these features in the context of gene-linked isoform sedimentation and found that shorter CDS, longer UTRs, lower 5’ UTR GC-content and higher 3’ UTR GC-content relative to the dominant isoform in the cytosol corresponded to monosomal and light polyribosomal sedimentation of isoforms. As a whole, our results present intrinsic feature measurements and potential sequence motifs that likely enact combinatorial effects on translation, providing both previously reported and novel insights into the underlying mechanisms of ASTC.

**Figure 5.**
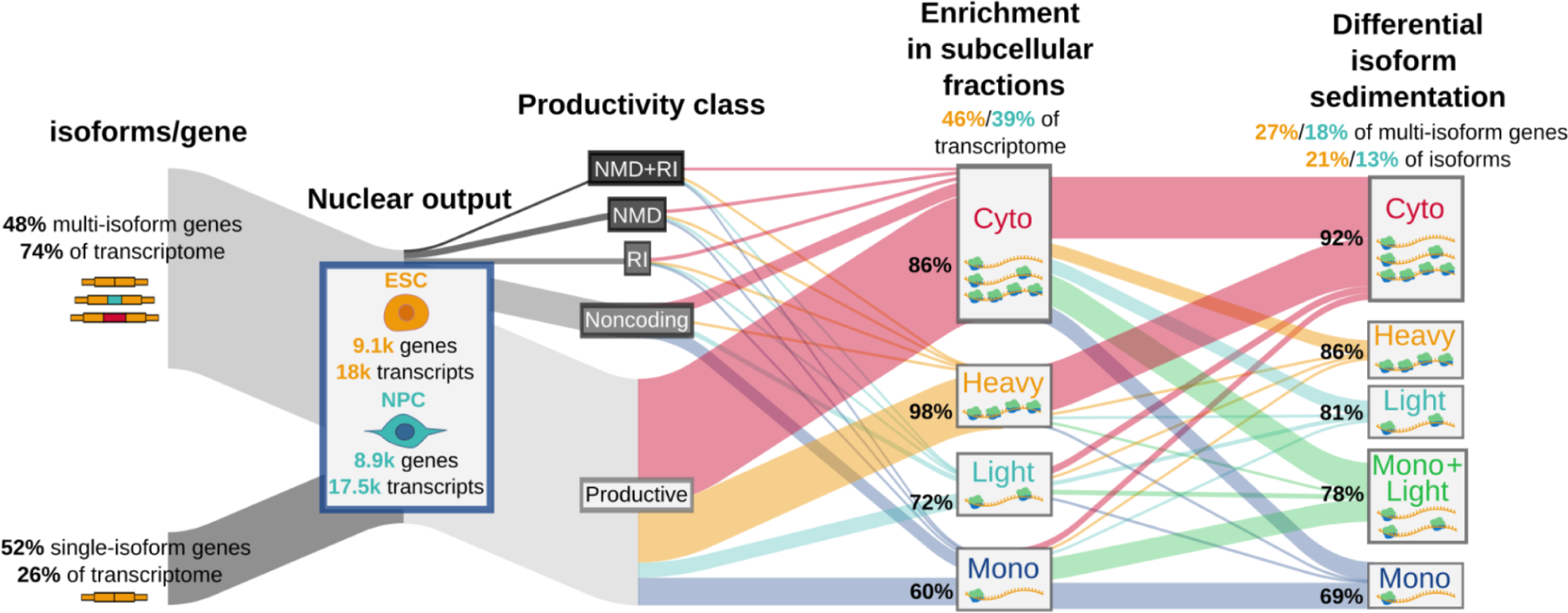
Summary of transcriptome characterization and ribosome association profiles. Transcripts were classified by the number of isoforms per gene they belong to and by their productivity. 46% and 39% of transcripts in ESC and NPC, respectively, were enriched in particular fractions. 27% and 18% of multi-isoform genes in ESC and NPC, respectively, exhibited differential isoform sedimentation. The proportion of productive transcripts in the context of fraction-associated enrichment and distinct isoform sedimentation are denoted as percentages to the left of each fraction.

Because the light and heavy polyribosome fractions were pooled sets of individual polyribosome fractions, we could not assess features in the context of ribosome density. To be clear, LR Frac-seq can be performed without pooling of individual polyribosome fractions, which would enable ribosome density-level analyses. We note that the heavy polyribosome fraction is likely composed of both efficiently and inefficiently translated transcripts depending on their ribosome density, and that trends of feature length and GC-content are subject to exceptions in each subcellular fraction. Additionally, Frac-seq differs from ribosome profiling methods in that it doesn’t capture single-nucleotide resolution ribosome footprints. Rather, it stratifies the translated transcriptome in terms of the number of ribosomes associated with full-length mRNAs. Therefore, it is not intended to replace ribosome profiling methods and is instead an alternative approach that benefits from retaining UTRs. We recommend LR Frac-seq for the study of translational control in cases where complete isoform structures and detection of novel isoforms is desired. For more complete coverage of the entire transcriptome, it may be beneficial to merge the LR transcriptome with an established, comprehensive transcriptome annotation for the organism of interest, like one available from GENCODE.

A major implication of LR Frac-seq in the field of translatomics is that its library preparation can be modified to enable direct RNA sequencing after fractionation to detect post-transcriptional modifications which are understood to significantly influence translation. For instance, RNA methylation, specifically N6-methyladenosine (m6A), can alter translation efficiency. Pseudouridylation can affect translation dynamics by influencing ribosome stalling and pausing during protein synthesis. RNA editing events, such as adenosine-to-inosine (A-to-I) editing, can modify regulatory sequences, altering the fate of mRNAs. These PTMs exemplify some of the multifaceted ways in which RNA modifications can impact translational control. By coupling accurate positions of post-transcriptional modifications with polyribosome profiles at isoform resolution, LR Frac-seq could enable more direct correlation of modifications with their effects on translation. Because we used R2C2, which is a cDNA method, to strengthen the confidence of isoform structures, we did not capture modification information beyond RNA editing events. But future adopters of LR Frac-seq can employ direct RNA sequencing methods after fractionation to gain that additional layer of data.

In conclusion, LR Frac-seq enables polyribosome profiling at isoform-level, retaining complete information about UTRs and novel transcript structures. We tested this method in the context of neuronal differentiation, revealing widespread enrichment of transcripts in subcellular fractions relative to the cytosol and largely congruent patterns of isoforms-specific sedimentation between ESC and NPC. Our results substantiate that intrinsic transcript features are an important determinant of ribosome association, and this work presents a promising new approach to study translational control without the information loss suffered by ribosome profiling and short read sequencing-based methods.

## METHODS

### H9 cell culture and differentiation to NPC

H9 cells in feeder-free culture were disaggregated using accutase and resuspended in hESC medium (StemMACS) containing 10 µM Rock inhibitor (Y27632). Cells were then seeded on a matrigel-coated 12-well plate at 50k live cells per well. Rock inhibitor was withdrawn the next day and the cells were cultured in hESC medium for 3 days. Neural differentiation was then induced over 7 days using KSR medium (for 500.5 mL stock: 415 mL KO-DMEM, 75 mL KSR, 100X Glutamax, 100X NEAA, 1000X bME, 10 µM SB431542, 100 nM LDN-193189). A subset of differentiated cells were stained for PAX6 to confirm neural differentiation.

### Short read Frac-seq

Cytosolic extracts from monolayer-cultured H9 cells and H9-derived NPCs, both in triplicate, were separated on sucrose gradients as described in the original Frac-seq publication (Sterne-Weiler et al. 2013). From these, the monosome fraction (RNAs associated with 1 ribosome), light polyribosome fraction (2-4 ribosomes) and heavy polyribosome fraction (≥5 ribosomes) were isolated using the Gradient Station (Biocomp Inc). RNA was extracted with TRIzol, polyA selected, and converted to directional RNA Seq libraries (BIOO Scientific qRNA) from these fractions in addition to total cytosolic RNA. Biological and technical replicates were sequenced using Hiseq 4000 PE150 (50-100M reads per library).

### Long read Frac-seq

From the same fractionated mRNA used prior for Illumina sequencing, full-length cDNA was prepared using the Rolling Circle Amplification to Concatemeric Consensus (R2C2) method (Volden et al. 2018). Libraries were pooled and sequenced on an ONT PromethION, generating 12.11M reads with read length N50 of 17.6Kb.

### *De novo* transcriptome assembly from long reads

R2C2 long reads were basecalled with Bonito v0.0.1 (https://github.com/nanoporetech/bonito). Subsequent polyA tail and adapter trimming followed by definition of high-confidence isoform consensus sequences was carried out using Mandalorion v4.0.0 (Volden et al. 2023) with all sample FASTAs (from ESC and NPC, all subcellular and cytosolic fractions in duplicate) as input. The resultant transcriptome was filtered for redundant transcripts using GFFCompare v0.12.6 (Pertea and Pertea 2020) against the GRCh38.p13 Release 41 primary assembly annotation (Harrow et al. 2012), and then further filtered and annotated using SQANTI3 v5.1.1 and IsoAnnot Lite v.2.7.3 (de la Fuente et al. 2020). SQANTI3 filtering was done using the machine learning filter with a training set proportion of 80% and a correct classification probability threshold of 70%.

### Short read data analysis

Short reads were adapter-trimmed with cutadapt, then mapped to the GRCh38.p13 primary assembly genome with the long read-derived transcriptome annotation using STAR v2.7.8a (Dobin et al. 2013). Transcript-level quantification was performed from the alignments using Salmon v1.9.0 (Patro et al. 2017) in alignment-based mode. Differential expression analysis at the gene, transcript, and isoform level were carried out using tappAS v1.0.7 (de la Fuente et al. 2020), which utilizes maSigPro v1.72.0 with the following analysis parameters: polynomial degree of 3, significance level of 0.05, R^2^ cutoff of 0.7, fold change of 2, and 9 K clusters. Differential expression analyses were performed for each subcellular fraction against its cognate cytosolic fraction (all in triplicate) for each cell type, and between subcellular and cytosolic fractions across ESC and NPC. Pathway analyses were done using Metascape (Zhou et al. 2019). Alternative splicing (AS) analysis was performed using junctionCounts (https://github.com/ajw2329/junctionCounts), which identifies and quantifies binary splicing events from RNA-seq data, including: alternative 5’ and 3’ splice sites (A5SS and A3SS), alternative first and last exons (AFE and ALE), skipped exons (SE), retained introns (RI), and mutually exclusive exons (MXE). AS events were then statistically tested by comparing the dispersions of junction support for their included and excluded forms using DEXSeq v1.46.0 (Anders, Reyes, and Huber 2012). Events were considered significant if they had 0.1 ≤ Ψ ≤ 0.9 and adjusted p-value ≤ 0.05 when assessing splicing within a condition, or |ΔΨ| ≥ 0.1 and Q-value ≤ 0.05 when assessing changes in splicing across conditions.

### Feature analysis

Transcript features were collected from the transcriptome IsoAnnot Lite annotation, including length measurements of: transcript, CDS, total upstream open reading frame (uORF), 5’ and 3’ UTRs. Total counts of: 5’ and 3’ UTR motifs, miR sites, uORFs and repeat sequences. Binary features: coding/noncoding, proximal/distal polyA tail usage, predicted nonsense-mediated decay (NMD)/no NMD and intron retention/no intron retention. And GC-content of the transcript, CDS, and 5’ and 3’ UTRs. Feature selection for binary classification between transcripts belonging to subcellular fractions was performed using the Random Forest Classifier method from the sklearn.ensemble module of scikit-learn v1.2.2 (https://scikit-learn.org/stable) and evaluated using permutation importance from the sklearn.inspection module.

Motif analysis was performed using HOMER v4.11 (Heinz et al. 2010). Target sequences were produced by slicing fraction-associated skipped exons (in the monosome relative to cytosol, the light polyribosome fraction relative to cytosol, and the heavy polyribosome relative to the monosome and the light polyribosome separately) into 30 nt windows; segregating the first 30 nt (start of the exon) and the last 30 nt (end of the exon) from the rest of the exon body (middle). Each group (start, middle, end) was subjected to *de novo* motif discovery against background sets of 30 nt windows produced from skipped exons that were not enriched in their given fraction. Significant motifs (p-value ≤ 0.05, FDR ≤ 0.2) were then tested for enrichment across all sets of windows in each fraction using SEA v5.5.4 (Bailey and Grant 2021). Enriched motifs with p-value ≤ 0.05 in at least one set of windows were then compared to RNA-binding protein motifs in the Ray 2013 Homo sapiens dataset (Ray et al. 2013) for potential matches using Tomtom v5.5.4 (Gupta et al. 2007). Position weight matrices for RNA-binding proteins with significant matches to motifs (p-value ≤ 0.05) were procured from CISBP-RNA (Ray et al. 2013) for visualization.

### Luciferase reporter assays

Luciferase reporters designed to test translational control by alternative first exon sequences were assembled from gene blocks (IDTDNA) and cloned into pLightSwitch 5’UTR report (Switchgear Genomics). HEK293 cells, grown on 6 well plates in DMEM supplemented with 10% FCS, were transfected with 2.5 µg pLightswitch reporter plasmid and pMIR (Ambion). 24 hours post-transfection, cells were lysed with Passive Lysis Buffer and analyzed by dual luciferase assay (Promega). Experiments were performed in triplicate. Relative luciferase activity (Renilla vs. Firefly) was plotted in Graphpad and analyzed by paired T-test.

## DATA ACCESS

All sequencing data are available through the Gene Expression Omnibus Short Read Archive (GSE244655).

## ACKNOWLEDGEMENTS

This research was made possible by a grant from the California Institute for Regenerative Medicine (GCIR-06673-A). The contents of this publication are solely the responsibility of the authors and do not necessarily represent the official views of CIRM or any other agency of the State of California. This work was also supported by grants from the National Institutes of Health R35GM130361.

## AUTHOR CONTRIBUTIONS

AJR performed data processing, bioinformatic analyses, and wrote the paper; JMD designed and performed experiments; JRS conceptualized the project and designed experiments.

